# MLL/WDR5 complex recruits KIF2C to midbody to ensure MT depolymerization and furrow compaction during cytokinesis

**DOI:** 10.1101/2024.10.17.618797

**Authors:** Avishek Kataria, Neeraja Hemalatha, Akash Chinchole, Shweta Tyagi

**Author notes:** Corresponding author: Laboratory of Cell Cycle Regulation, CDFD Uppal, Hyderabad 500039, India.

## Abstract

Mixed-lineage leukemia (MLL) protein is a well characterized epigenetic regulator whose non-canonical activities remain under appreciated. Here we show that MLL and its associated protein WDR5, localize to the midbody. Loss of MLL/WDR5 results in defective midbody formation, which displays a wide midzone-like microtubule structure, along with chromosome bridges, resulting in binucleated cells. We show that MLL and WDR5 interact with kinesin 13 motor—KIF2C, and targets it to the midbody. The depolymerase activity of KIF2C promotes correct localization of centralspindlin complex, compaction of midzone MTs and finally furrow completion. By characterizing the previously undiscovered role of MLL and KIF2C in the regulation of cytokinesis, our work underscores the importance of these proteins at the interface of actin and microtubule cytoskeleton regulation, pathways that are frequently altered in oncogenesis.

## Introduction

Mixed-lineage leukemia (MLL/KMT2A) protein, frequently mutated or translocated in cancer, is the founding member of evolutionary conserved lysine methyltransferases KMT2 family, responsible for depositing the mono-, di-, tri-methylation marks on Histone 3 Lysine 4 (H3K4). Interestingly, this KMT2-family enzyme requires the WRAD complex, composed of WDR5, RbBP5, Ash2L and Dpy30 proteins, to enhance its SET-domain mediated HMT activity (Patel et al., 2008; Song & Kingston, 2008; Sugeedha et al., 2021).

MLL has been well studied as an epigenetic transcriptional regulator. However, its epigenetic contributions in cancer have been questioned (Cao et al., 2014; Mishra et al., 2014). Previously, we have identified various mitotic defects associated with loss of MLL complex, including defects in chromosome alignment, spindle assembly and cytokinesis aberrations, which may lead to micronuclei formation or binucleation, thus promoting genomic instability (Ali et al., 2014, 2017). Interestingly MLL does not utilize its SET domain-mediated methyltransferase or TAD domain-mediated transcriptional activity for this function. Instead, MLL’s interaction with WDR5 is essential for these activities (Ali et al., 2014). Further, we have shown that MLL localizes to centrosome, spindle apparatus and midbody during mitosis (Karole et al., 2018). Our studies have revealed that MLL/WDR5 complex plays a crucial roles on the spindle apparatus and centrosome to ensure proper spindle assembly (Ali et al., 2017; Chodisetty et al., 2024). However, whether MLL/ WDR5 complex plays a critical role in the formation of midbody and in the process of cytokinesis, still remains undiscovered.

Cytokinesis, the final step in cell division, ensures that dividing cells are partitioned correctly into two daughter cells. Aberrations in cytokinesis are associated with genetically unstable cells, aneuploidy and oncogenesis (Li et al., 2018; Petsalaki & Zachos, 2021). During the process of cytokinesis, the bipolar microtubule (MT) forms an array between the separating sister chromatids, also called the midzone or central spindle, which gives rise to the midbody. The transformation of midzone into midbody is linked to furrow ingression; blocking this process leads to the accumulation of midzone-like MT structures (Straight et al., 2003). The furrow compacts the antiparallel midzone MT into a bundle, with a central bulge known as the stem body emerging during compaction. Several models explaining role of astral or central spindle MT in furrow ingression have been proposed where MT-mediated delivery of their associated proteins is considered a critical event to bring about a signal cascade regulating the furrow formation from the cortex (D’Avino et al., 2005). Interestingly MT dynamics is required for furrow initiation and completion (Shannon et al., 2005). Here we describe the role of MLL/WDR5 complex in targeting the Mitotic centromere associated kinesin (MCAK)/KIF2C, a motor protein from Kinesin-13 family, to the midbody. Like other members of this family, KIF2C contains a kinesin motor domain and plays an important role in MT depolymerization and hence dynamics (Ems-McClung & Walczak, 2010; Ritter et al., 2015; Steblyanko et al., 2020). We show that MLL and WDR5 localize to the midbody and their loss results in aberrant midbody formation. MLL/WDR5 interact with KIF2C and target it to the midbody to bring about MT depolymerization. KIF2C mutant, defective for MT depolymerization, phenocopies the aberrant midbody phenotype and exhibits defective furrow compaction, indicating that role of KIF2C extends beyond metaphase.

## Results and Discussion

### MLL and WDR5 promote the formation of the midbody

In our earlier work, we have reported that MLL/WDR5 complex localize to spindle apparatus (Ali et al., 2017). Additionally, we have observed that MLL can also be found on mitotic chromatin, centrosomes, central spindle, and midbody at different stages of mitosis (Karole et al., 2018). Indeed, loss of MLL or WDR5 gave rise to binucleated cells indicating defects in cytokinesis (Ali et al., 2014; Bailey et al., 2015). Previously, we have optimized different fixation conditions to observe MLL localization on individual mitotic structure including midbody (Ali et al., 2017; Karole et al., 2018). Upon immunofluorescence staining (IFS), initially MLL appeared as two closely space bands, characteristic of dark zone proteins (Fig 1A; (Hu et al., 2012)). However, later with the formation of the stem body, we could observe it as a single band characteristic of bulge/stem body proteins (compare MLL staining in panel a to panel b, Fig 1A). In fact, MLL could be visualized as a single ring by structured illumination microscopy (Fig 1B, Supp Fig S1A). In contrast to MLL, we observed that WDR5 appeared as four bands, inner two bands characteristic of dark-zone proteins (Fig 1C, red arrows), whereas the outer bands likely corresponded to flanking-zone proteins (Fig 1C, white arrows). MLL and WDR5 typically show high degree of co-localization. After further analysis, we could observe WDR5 on the stem body, between two outer bands in some images (Fig 1D). Most likely WDR5 also localizes on the stem body like MLL, but its antibody staining is blocked in other images, as is characteristic of this zone.

**Figure 1:**
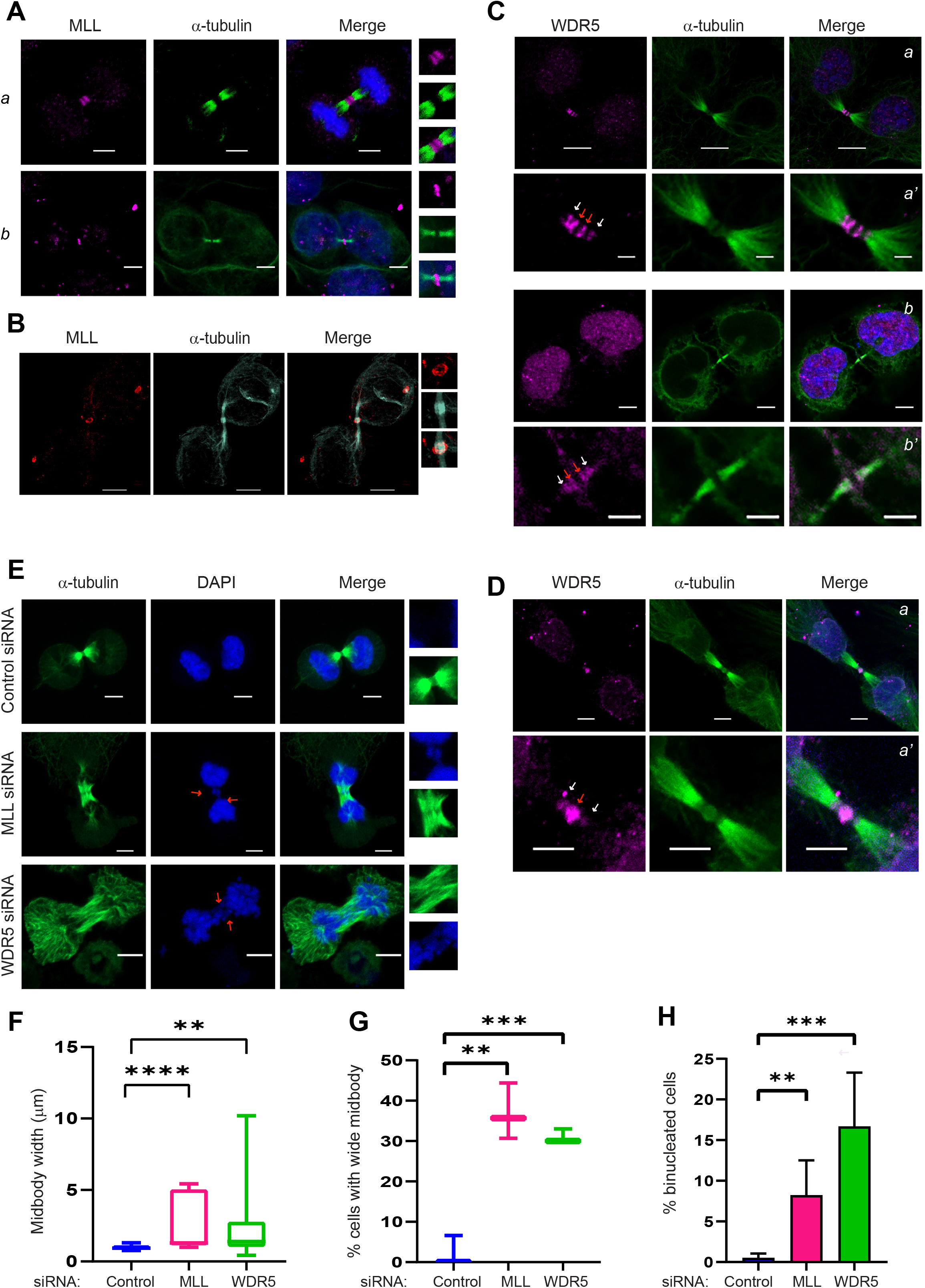
Loss of MLL/WDR5 affects the midbody formation. (**A-D**) U-2OS cells were fixed and used for IFS staining (IFS). Cells were stained with MLL (**A, B**), WDR5 (**C, D**), and α-tubulin (**A-D**) antibodies. Condensed chromosomes were stained with 4’, 6’-diamidino2-phenylindole (DAPI). Zoomed in images are shown inset. (**B**) Images were acquired in 3D-structured-illumination microscope (3D-SIM) as shown. (**C, D**) Images show WDR5 localization at midbody. a’ and b’ are magnified images of panel a and b respectively. (**E**) U-2OS cells, treated with control, MLL, or WDR5 siRNA for 72 h were fixed and used for IFS. Red arrows indicate lagging chromosomes. (**A-E**) Scale bars: 5 μm. (**F-G**) Quantification of (**F**) midbody width, and (**G**) percentage of cells with wide midbody, as displayed in **E**, from three independent experiments, is shown. (**H**) Percentage of binucleated cells in MLL and WDR5 siRNA were quantified from three independent experiments (see also supplemental Fig S1F-G). Data represents Mean ± S.D., unpaired Student’s t-test was used for comparing control vs siRNA condition. \*\*\*\**P*<0.0001,\*\*\**P*<0.001, \*\**P*<0.01.

In order to confirm specific localization of our proteins, we performed RNAi. Protein levels of both MLL and WDR5 decreased on siRNA treatment and this could be observed both in the immunoblot (Fig S1B, C) as well as immunofluorescence staining (IFS) (Fig SF1D, E), indicating that our IFS of MLL and WDR5 was specific. During RNAi experiments, we observed that a significant proportion of MLL or WDR5 siRNA-treated cells displayed a wide midbody (Fig 1E). In these cells, it seemed as if conversion of midzones MTs had started but were either delayed or defective to form a midbody (Figure 1E; Hu et al., 2012). When we measured the width of this incomplete midbody (hereafter referred to as In-midbody), it was about 2 times or more than those observed in control cells, and was displayed by about 40% MLL or WDR5 siRNA-treated cells (Fig 1F-G). The In-midbody was strikingly enriched in MTs and displayed a dense array when compared to control siRNA-treated cells (Fig 1E). We also observed chromosome bridges in these cells indicating that loss of our proteins did not cause a mere delay in the process but had a more vital role in the chromosome segregation and midbody formation. Consistent with our claim above and our previous results, we observed significant number of binucleated cells in MLL or WDR5 siRNA-treated cells, indicating that MLL/WDR5 complex was involved in midbody formation and ensuing cytokinesis (Fig 1H, S1F-G) (Ali et al., 2014).

### MLL and WDR5 interact with Kinesin 13 family members —KIF2C and KIF2B

In our previous studies, based on extensive complementation experiments using MLL and WDR5 mutants, we have reported that defects in cytokinesis are independent of MLL’s transcriptional activity, and are dependent on protein-protein interactions of MLL with WDR5 and its partnering proteins (Ali et al., 2014, 2017). At this time we also reported that WDR5 interacts with all three members of kinesin 13 motors, namely KIF2A, KIF2B and KIF2C, and characterized the interaction of MLL-WDR5-KIF2A in details showing the role of MLL/WDR5 in targeting KIF2A to the spindle poles (Ali et al., 2017). The dense MT staining observed here is reminiscent of the deregulated MT depolymerase activity of a kinesin 13 motors (Ali et al., 2017). While KIF2A has been reported to localize to central spindle, phosphorylation by Aurora B limits KIF2A to distal ends of the central spindle, thereby excluding it from the spindle midzone (Ganem & Compton, 2004; Uehara et al., 2013). In contrast, KIF2B and KIF2C have been reported to be on the midbody (Manning et al., 2007; Qin et al., 2016; Zhang et al., 2011) and detected in midbody interactome (Capalbo et al., 2019). Therefore, we decided to probe for interaction of KIF2B and KIF2C with the MLL complex proteins. We used bacterially expressed GST or GST-WDR5 fusion protein to perform pull-down experiments. As the antibodies for KIF2B and KIF2C are not available readily, we made stable cell lines expressing GFP fusions of these proteins. The cell lysates from GFP-cell lines were used for pull down experiments. As shown in Fig. 2A GST-WDR5 could readily pull down KIF2B and KIF2C while the control (GST) was clean. Next, we performed reciprocal pull down using GST-KIF2C (and GST-KIF2B) and found them interacting with WDR5 and RbBP5, indicating that MLL complex proteins associated with these kinesins (Fig 2B, S2A).

**Figure 2:**
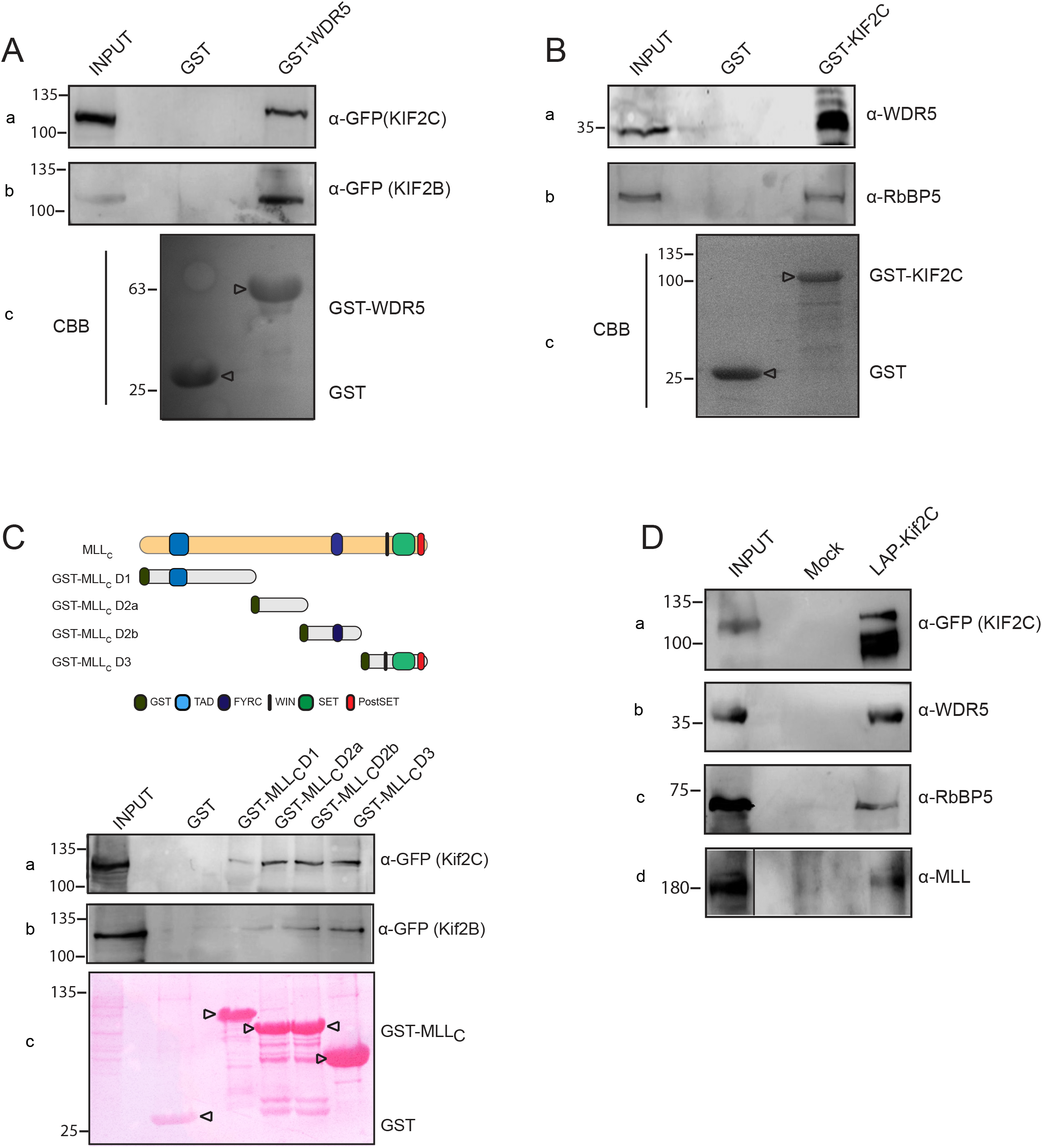
MLL and WDR5 interact with kinesin 13 motors—KIF2C and KIF2B. (**A**) The HEK293T cells `stably` expressing `GFP-`KIF2B and `GFP-`KIF2C were lysed and `the` extract was subjected to affinity pull-down using GST `or` GST-WDR5 bound beads. The immunoblot was probed with GFP to detect GFP-KIF2B and GFP-KIF2C proteins. Amount of GST protein used for the assay is shown using shown by Coomassie brilliant blue (CBB) staining (panel c). Relevant bands are indicated using arrow head. (**B**) The HEK293T cell lysate was subjected to affinity pull-down using GST, or GST-KIF2C bound beads. The immunoblot was probed with anti-WDR5 and anti-RBBP5 antibody to detect endogenous proteins. (**C**) Schematic of MLLc subunit and various GST-tagged MLLc deletion constructs are shown. The lysate from HEK293T cells expressing GFP-KIF2B and GFP-KIF2C was used for affinity pull-down using GST, GST-MLLCD1, GST-MLLCD2a, GST-MLLCD2b and GST-MLLCD3 bound beads. The immunoblot was probed by using GFP antibody. Amount of GST proteins used for pull-down assay is shown using Ponceau S staining. (**D**) HeLa Kyoto cell lines expressing LAP Tagged KIF2C.GFP were lysed and affinity pulled down using S-protein beads. Immunoblots were stained using anti-GFP, anti-WDR5, anti-RbBP5 and anti-MLL antibody. (**A-D**)Numbers on the left indicate the position of molecular weight markers (in kDa).

MLL C subunit is a large protein. In order to interrogate if these kinesins interacted with it and to map the region of interaction, we created 4 truncations of MLLC protein as shown in Fig. 2C. Using our U-2OS stable cell lines expressing GFP-KIF2B and GFP-KIF2C we found that these proteins interacted with most of MLLC including the FYRC and SET domain (Fig 2C). It is important to note here that KIF2A also interacted with the same regions of MLLC [Note: Due to unstable and difficult expression, we divided the MLLC protein into four parts (instead of three) such that D1 and D2 gave rise to three deletions and D3 remained unchanged; (Ali et al., 2017)], indicating that the kinesin 13 motors exhibited similarity in their interaction with MLLC subunit.

Finally, we made use of localisation and affinity purification (LAP) Hela cell lines expressing KIF2B or KIF2C from a BAC which drives the expression of promoter for respective protein. The tag provides fusion with S-protein for affinity purification, and GFP for localisation (Maliga et al., 2013; Poser et al., 2008). We perform pull-down experiments using S-protein beads and checked their interaction with endogenous MLL complex proteins as shown in Fig 2D. WDR5, RbBP5 and MLL could be readily pulled down by these motors indicating that just like KIF2A, KIF2B and KIF2C also occurred in a complex with MLL core components (Fig 2D, S2B, C).

### MLL/WDR5 complex targets KIF2C to the midbody

Given that both KIF2B and KIF2C interacted with the MLL complex and have been reported to localize at the midbody, we performed IFS to confirm their midbody localization. We could see KIF2C accumulating on the midzone in anaphase though it primarily stained the centromere/kinetochores and spindle poles (Fig 3A, panel a). However, during telophase and later, KIF2C appeared as two closely spaced dark-zone bands, which also showed up in the stem body in some images (Fig 3A, panel b, 3F, S3A panel b). The same observation was made for KIF2B during telophase, where it showed up as dark-zone protein first and on the stem body subsequently (Fig S3A).

**Figure 3:**
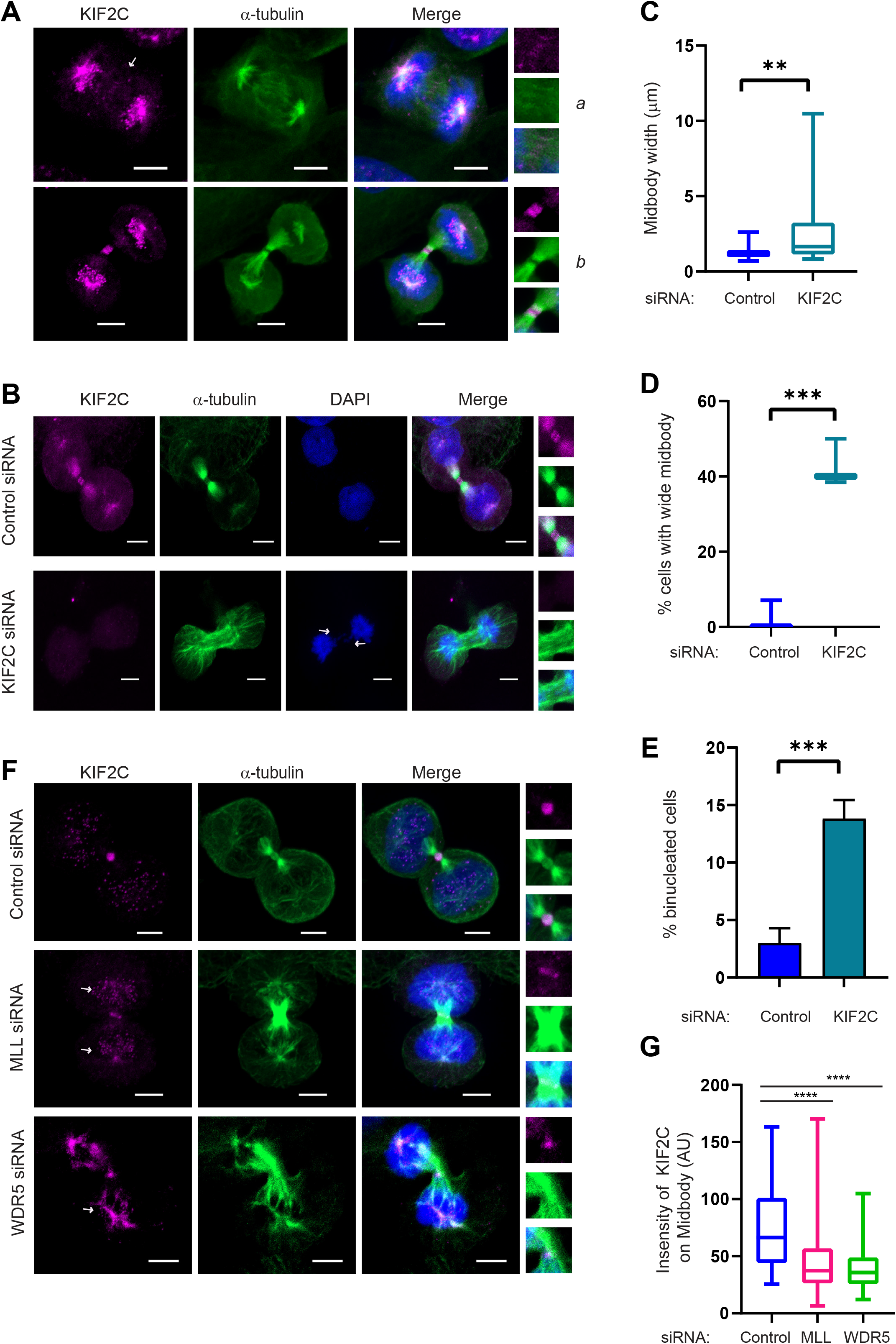
MLL/WDR5 complex targets KIF2C to the midbody. (**A**) IFS of U-2OS cells stained with KIF2C antibody. White arrow indicates KIF2C on midzone. (**B**) After treatment with control and KIF2C siRNA for 72 h, U-2OS cells were fixed and stained with antibodies against KIF2C, and α-tubulin. White arrows indicate lagging chromosomes. Quantification of (**C**) midbody width, and (**D**) percentage of cells with wide midbody, as displayed in **B**, from three independent experiments, is shown. (**E**) Percentage of binucleated cells in KIF2C siRNA were quantified from three independent experiments (see also supplemental Fig S3C). (**F, G**) U-2OS cells were treated with control, MLL or WDR5 siRNA for 72 h. Cells stained with KIF2C and α-tubulin antibody were quantified for intensity of KIF2C at midbody (**G**) and centromeres (**S3D**)(white arrows indicate KIF2C localization at centromere). Data represents Mean ± S.D., unpaired student T-test was used for comparing control vs siRNA condition. \*\*\*\**P*<0.0001, \*\*\**P*<0.001, \*\**P*<0.01. Scale bars: 5 μm.

At this time we were able to procure a working antibody for KIF2C (but not KIF2B) limiting our further experiments to KIF2C. We depleted KIF2C with previously published siRNA. The protein levels showed substantial decrease in Western blot (Fig S3B) and IFS (Fig 3B). Pleasantly, like MLL and WDR5 siRNA treated cell, 40% KIF2C siRNA treated cells also displayed an In-midbody phenotype with the midbody width significantly more than the control cells (Fig 3B-D). The cells with In-midbody phenotype displayed chromosome bridges and dense MTs. We also observed high number of binucleated cells in KIF2C-siRNA treated cells (Fig 3E).

Next, we asked if MLL and/or WDR5 target KIF2C to the midbody. We treated the cells with MLL or WDR5 siRNA and stained them for KIF2C and α-tubulin (Fig 3F). In MLL-depleted cells, KIF2C showed substantial decrease in midbody-staining. Remarkably this decrease in KIF2C intensity was not observed on the centromeres suggesting that MLL had specific association/function with KIF2C (Fig 3F, G, S3D). The same observation was made in WDR5-depleted cells indicating that MLL and WDR5 acted together to bring KIF2C to the midbody (Fig 3F, G, S3E).

### The MT depolymerase activity of KIF2C is required for furrow ingression during cytokinesis

The images with loss of KIF2C suggest that the In-midbody phenotype is most likely due to decrease in MT depolymerization during midzone to midbody transition brought about by KIF2C and it is phenocopied by MLL/WDR5 as they recruit KIF2C to the midbody. MT dynamics as well as ordered assembly and disassembly is important for the formation of the midbody (Elia et al., 2011; Hu et al., 2012; Straight et al., 2003). In order to prove that the depolymerase activity of KIF2C is involved, we used siRNA-resistant constructs of KIF2C expressing either the wild type or mutant (G495A) protein defective for ATP hydrolysis and therefore MT depolymerization (Wang et al., 2012, 2015) to generate stable cell lines. Upon siRNA-treatment, the cells expressing wild type KIF2C formed midbody like the Control cells (Fig 4A, S4A), while the KIF2C G495A mutant behaved like U-2OS cells (alone) depleted of KIF2C. We observed a wide midbody along with dense MT as well as chromosome bridges (Fig 4A). When measured the midbody width was high in U-2OS cells alone or expressing the KIF2C mutant but not in cells expressing wild type KIF2C (Fig 4B).

**Figure 4:**
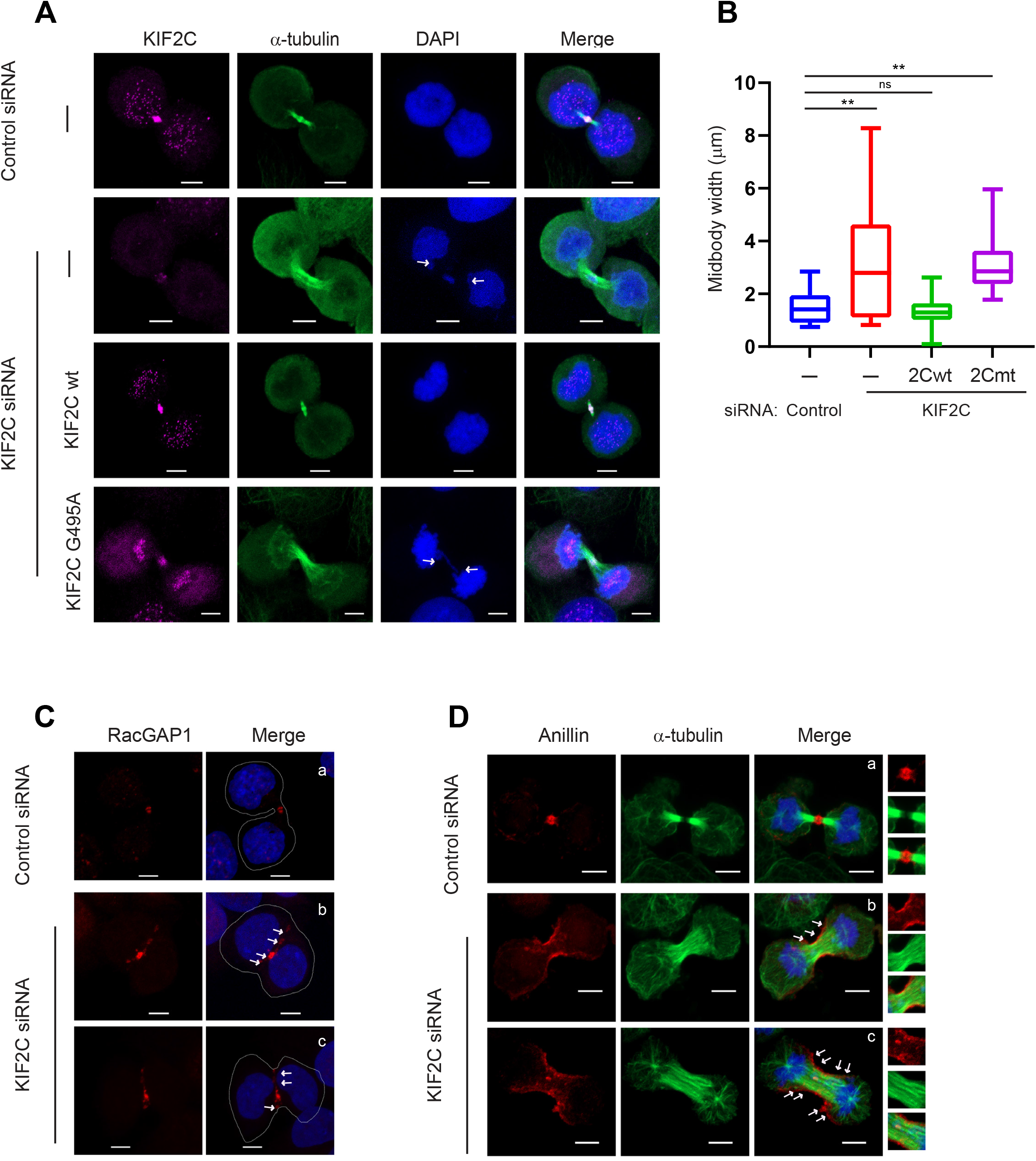
MT depolymerization activity KIF2C plays role in cytokinesis. (**A, B**) Wild-type U-2OS cells and cells expressing siRNA resistant GFP-KIF2C and GFP-KIF2C G495A were treated with KIF2C siRNA to deplete the endogenous KIF2C, and stained with KIF2C, and α-tubulin. Midbody width was measured as shown in B. Data represents Mean ± SD. \*\**P*<0.01, ns: not significant (Mann-whitney unpaired t-test). (C, D) U-2OS cells were treated with Control vs KIF2C siRNA for 72 h. Cells were fixed and stained with indicated antibody – RacGap1(**C**), Anillin & α-tubulin (**D**). Scale bars: 5 μm.

MTs are also essential for cleavage furrow positioning and stem body alignment during cytokinesis (Shannon et al., 2005; Uehara et al., 2013). RacGAP1, a component of centralspindlin complex, is essential for cleavage furrow ingression (Simon et al., 2008). To determine what may be happening in absence of KIF2C, we performed IFS with anti-RacGAP1 antibody. We observed that in Control cells, RacGAP1accumulated into a single cluster at the intercellular bridge (Fig 4C). However, KIF2C-depleted cells displayed scattered signals of RacGAP1, along the equatorial plane, indicating that proper central spindle organization could not be attained in these cells. This phenotype has been also reported in cells with Taxol stabilized MTs (Uehara et al., 2013). While drawing the outline of the KIF2C-siRNA treated cells, we observed that the furrow ingression was also retarded in these cells (Fig 4C). In order to confirm this observations, we used contractile ring protein Anillin to determine the progress of furrow ingression (Fig S4B). As shown in Fig 4D, unlike the Control cells, the KIF2C-siRNA treated cells displayed Anillin still at furrow cortex, presumably unable to successfully progress to the contractile ring constriction. Our findings here indicate that the MLL/WDR5-targeted MT depolymerase activity of KIF2C is required to maintain MT dynamics for successful re-localization of microtubule-binding proteins involved in compaction of midzone MTs and furrow completion.

Here we show for the first time that KIF2C, recruited by MLL/WDR5 complex, has a function beyond correcting erroneous attachments of microtubule-kinetochore and spindle formation. Our results demonstrate that KIF2C stains as the dark zone protein on the midbody and its activities as a MT depolymerase are responsible for the successful migration of central spindle complex and furrow compaction.

Here we have shown that MLL regulates the midbody formation by regulating spindle dynamics via KIF2C depolymerase activity. Remarkably, both MLL and KIF2C have been linked to the actin dynamics before. MLL may be involved in regulating the actin dynamics during the contractile ring formation by its regulatory role in the homeostasis of RhoA protein (Chinchole et al., 2022); while KIF2C has been shown to affect the actin-MT cytoskeleton dynamics thereby modulating cell migration (Moon et al., 2021). Interestingly, KIF2C is recruited by MLL/WDR5 complex selectively to the midbody but not the centromeres. We had made the same observation with KIF2A earlier as its localization was abrogated on the spindle poles, but not on centromeres upon, loss of MLL. We have recently shown that MLL regulates the transcription of centromeric RNA using its HMT activity (Malik et al., 2023). It is likely that the chromatin bound-MLL exists in different complexes/pools at the centromere than the KIF2A/C bound MLL complexes on the MT rich structures like midbody. In any case, our work here is adding another dimension to how MLL regulates genomic stability and why mutations in this protein are associated with a variety of cancers.

## Materials and Methods

### Cell Culture

Different cell lines used in our experiments are - U-2OS (Human osteosarcoma), HeLa (Human cervical adenocarcinoma) and HEK293T (Human embryonic Kidney cells). Dulbecco’s modified Eagle’s medium (DMEM, HiMedia, AT186) was used to culture different cell lines. 10% fetal bovine serum, L-glutamine and penicillin/streptomycin were added to DMEM before use. U-2OS cells stably expressing GFP-tagged KIF2C, KIF2B and KIF2C G495A were obtained by transfecting cells with appropriate DNA construct using polyethylene imine (PEI; Polysciences, 23966-2). HeLa kyoto cells stably expressing LAP tagged KIF2B, and KIF2C were a kind gift from A. Hyman (Maliga et al., 2013; Poser et al., 2008).

### Cloning

GST-WDR5, GST-KIF2B and GST-KIF2C constructs were described previously (Ali et. al 2017). pGEX4t1 was linearized with XhoI, and PCR-amplified MLL fragments were cloned to generate GST-MLLCD1(2723-3092aa), GST-MLLCD2a(3084-3445aa), GST-MLLCD2b(3448-3694aa), GST-MLLCD3(3694-3969aa). EGFP-KIF2B (#29479, kind gift from L. Wordeman), mEmerald-KIF2C (#54161, kind gift from M. Davidson, referred to as GFP-KIF2C throughout) cDNAs were obtained from Addgene. KIF2C was made siRNA resistant and in this construct G495A mutation was introduced using site-directed mutagenesis (Fig S4A).

### siRNA transfections

siRNA Transfections were done using Oligofectamine (Thermo Fisher scientific) as described previously (Tyagi and Herr 2009). After 72 hours of siRNA treatment, cells were harvested and used accordingly, either they were lysed in SDS laemelli buffer for immunoblotting experiment or they were fixed using different fixatives for microscopy experiments. Information about control, MLL and WDR5 siRNA have been published previously (Ali et al., 2014). Oligonucleotide 5’AACTCTAGGACTTGCATGATT 3’ was used to target KIF2C.

### Immunoblotting

Different cells used for immunoblotting were lysed using 2X NETN lysis buffer (40 mM Tris-Cl (pH 8.0), 200 mM NaCl, 1 mM EDTA, and 1% NP-40) supplemented with a freshly prepared protease inhibitor cocktail. After lysis, the laemelli buffer was added and the samples were boiled for 10 min. we used bradford assay to determine the concentration of protein in different samples. Different samples having equal amount of protein were loaded and resolved by SDS-PAGE. Using western blotting, proteins were transferred to surface of PVDF or Nitrocellulose membrane. For immunoblotting, membranes were incubated with following antibodies - MLL, KIF2C, WDR5, α-tubulin, RbBP5 and GFP. Details of different antibodies are provided below. Appropriate secondary antibodies were used for probing the blot, blots were developed using Amersham ECL substrate.

### Pulldown and Affinity purification

Various GST and GST tagged proteins (WDR5, MLLC D1, D2a, D2b, D3, KIF2B, and KIF2C) were produced in BL21 (DE3) pLysE bacterial cultures. After induction and protein production, bacterial cells were lysed using lysis buffer (50mM Tris pH 7.5, 150 mM NaCl, 1mM DTT, 0.5mM PMSF, 0.02% NP-40,). Glutathione Agarose beads (Sigma-Aldrich, G4510) were used to affinity purify the GST proteins from this lysate.

Mammalian cell lines HeLa, HEK-293T, U-2OS or stable cell lines were lysed using NETN buffer (100mM NaCl, 20mM Tris pH 8.0, 0.5mM EDTA, 0.2% NP-40, protease inhibitors). The cells were then lysed in NETN cell lysis buffer at 4°C on a rotator for 60 min. After lysis samples were centrifuged at 160003g for 15 minutes. After centrifugation the supernatant is collected and used in downstream process for pull down experiments.

GST Bound beads were added to the cell lysate and pull-down experiment was performed overnight at 4C on a rotator. Following pull-down, the GST bound beads were collected by centrifugation and washed 3 times with Wash buffer (50 mM Tris-HCl pH 7.4, 100 mM KCl, 0.05% Nonidet P-40). After washing, GST beads carrying the bound proteins were boiled in SDS laemelli buffer and used for immunoblotting.

S-protein coupled to agarose beads (Novagen) were used to affinity purify the LAP-tagged proteins from HeLa cell lysate. For pull-down, Hela Cell lysate was incubated with agarose beads overnight at 4C on a rotator. Following pull-down, the S protein beads were washed with Wash buffer and boiled in Laemmli buffer. The samples were resolved using SDS-PAGE and subsequently used for immunoblotting.

### Immunofluorescence staining

Different Cells used in our immunofluorescence assays were grown on coverslips. For arresting cells in G2/M phase, cells were treated with nocodazole (100 ng/ml) for 12 to 16 h. After Nocodazole arrest, cells were released into fresh medium for different time periods as mentioned in figure legends. Two different fixatives were used for staining different proteins - Methanol and paraformaldehyde. We used 4% paraformaldehyde for staining KIF2C, MLL and WDR5, cells were treated with fixative for 10min at room temperature. After fixation, permeabilization of the cells were done using 0.2% TX-100. Differently cells were also fixed with methanol for different duration for WDR5 (3 min) and MLL (6 min). After fixation cells were incubated with appropriate primary and secondary antibodies as described previously (Ali et al. 2017). DAPI (4’-6-diamidino-2-phenylindole) was used to label the chromatin following which coverslips were mounted using VECTASHIELD mounting medium (Vector laboratories-H1200). ZEISS LSM900 inverted confocal microscope and ZEISS Elyra inverted microscopes were used for imaging the cells as mentioned in figure legends. For SIM acquisition, the images were processed using ZEN 3.0 SR software.

While acquiring Z-stack images, a step size of 0.5 um was used. For quantification z-stacks were used to generate a maximum intensity projection (MIP) image. To quantify Midbody signal of MLL, WDR5 and KIF2C, a boundary was drawn manually around midbody in MIP images and mean intensity values for different channels of that region were calculated. Midbody width was calculated manually by using the distance tool present in Zen software. Centromere intensity was measured by placing a circle around the centromere foci of uniform diameter around all the centromere and mean intensity value for each channel was noted. To remove the background signal, a circle of equal diameter was placed adjacent to centromere. Mean intensity value of background signals were subtracted from centromere signal to get centromere intensity of the protein. Number of binucleated cells were counted manually. Various signal intensity measurements were carried out using Zen desk ver3.4.91 software.

### Antibodies

Various antibodies used in our experiments are as follows: Anti-WDR5 (Bethyl, A302-430A), Anti-MLLc (Bethyl, A300-374A), Anti-MLLc D1 (Chinchole et al., 2022), anti-GFP (Invitrogen, A11122), anti-RbBP5 (Bethyl, A300-109A), anti-KIF2C (protein-tech, 28372-1-AP), anti-α-tubulin (Sigma-Aldrich, T1568), anti-RACGAP1 (Cloud-Clone corp., PAE293Hu01), anti-Annilin (Cloud-clone corp., PAJ640Hu01), Alexa fluor 488 (Invitrogen, A11029, A11034), Alexa fluor 594 (Invitrogen, A11032, A11037, A21029) and Horse radish peroxidase (HRP) conjugated secondary antibodies (Bio-Rad, 170-6516, Bio-Rad, 170-6515).

### Statistical analysis

We used Graphpad Prism 9.3 for our quantitative and statistical analysis. Different tests applied on experiment data such as Unpaired student’s t-test and Mann-Whitney unpaired t-test are mentioned in legends along with their significance. Error bars in figures represent standard deviation (SD). Information about technical or biological replicates of experiments is provided in the figure legends. Each experiment was repeated minimally three times.

## Acknowledgements

We thank L. Wordeman and M. Davidson, for cDNA constructs; A. Hyman for HeLa LAP cell lines; A.K. is recipient of Junior and Senior Research Fellowships from CSIR, India towards the pursuit of a Ph.D. degree of the Manipal Academy of Higher Education. This work was supported in part by the DBT/Wellcome Trust India Alliance Senior Fellowship to S.T.[IA/S/18/2/503981] and CDFD core funds.

## Author contributions

A.K. performed most of the experiments. N.H. performed experiments related to Figures 2 and S2 and A.C. performed experiments related to Figures 1A-D, SF1D, E, G. S.T. and A.K. analyzed the results and wrote the manuscript.

