## Supplemental Information for "MLL/WDR5 complex recruits KIF2C to midbody to ensure MT depolymerization and furrow compaction during cytokinesis"

### Supplementary Information

**Figure S1**: **Loss of MLL and WDR5 leads to binucleated cells**. (A) U-2OS cells, fixed using methanol and stained with MLL antibody, were imaged in 3D-structured-illumination microscope (3D-SIM) as shown. (**B,C**) U-2OS cells, treated with control and MLL (**B**) or WDR5(**C**) siRNA for 72 h, were used for immunoblotting. Blots were probed with MLL, WDR5, KIF2C and α-tubulin antibody as shown. (**D-E**) Cells were fixed and stained with indicated antibody MLLc (**D**) and WDR5 (**E**) and used for IFS after siRNA treatment as shown. (**F-G**) Control, MLL (**F**) or WDR5 (**G**) siRNA treated cells were scored for binucleated cells after being stained with α-tubulin as shown. Scale bars: 5 μm (**A,D-E**) and 20 μm (**F-G**).

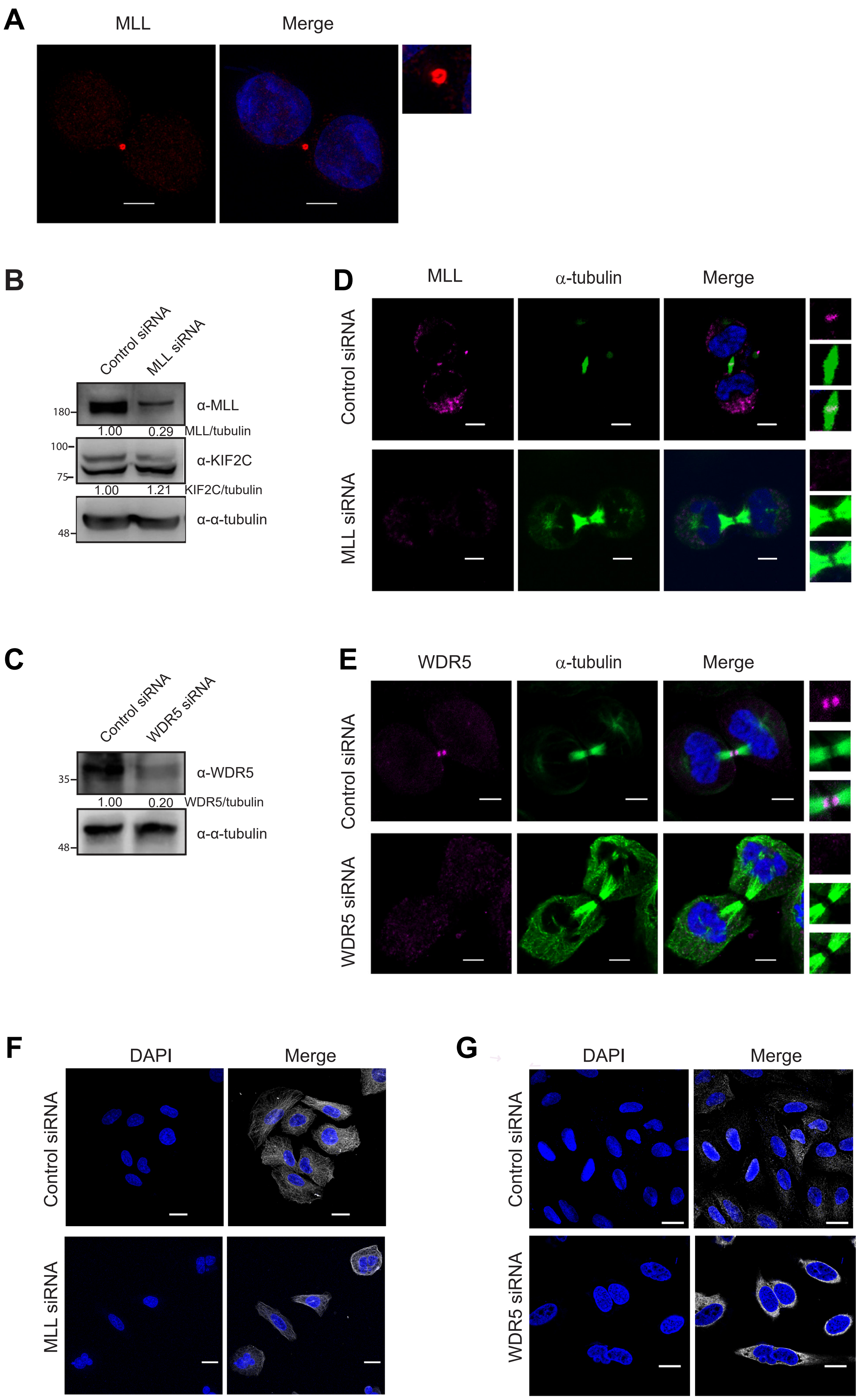


Supplementary Figure 1

**Figure S2: KIF2B interacts with MLL complex proteins.** (**A**) The HEK293T cell lysate was subjected to affinity pull-down using GST, and GST-KIF2B bound beads. The immunoblot was probed with anti-WDR5 and anti-RBBP5 antibody to detect endogenous proteins. Amount of GST protein used for the assay is shown by Coomassie brilliant blue (CBB) staining (panel c). Relevant band are indicated using arrow head. (**B-C**) HeLa cell lines expressing LAP Tagged KIF2B.GFP were lysed and subjected to affinity pulled down using S-protein beads. Immunoblots were stained using anti-GFP, WDR5 and RbBP5 (**B**); anti-GFP and MLL (**C**) antibody.

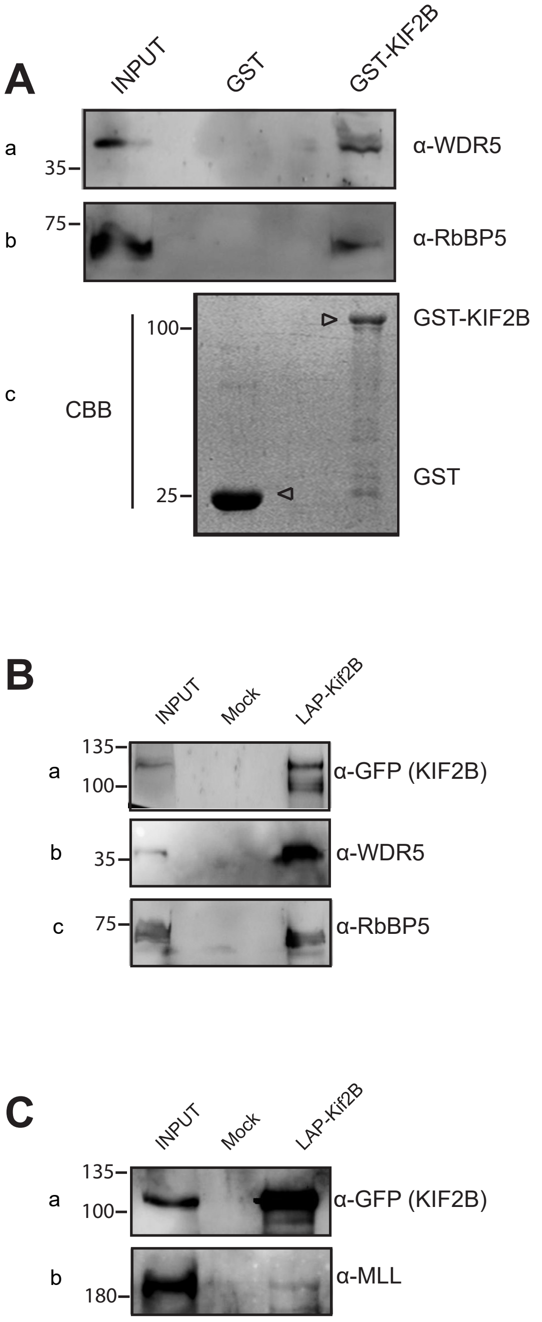


Supplementary Figure 2

**Figure S3**: **Loss of KIF2C affects cytokinesis.** (**A**) U-2OS cells expressing GFP-KIF2B were fixed using methanol and stained with different antibodies - GFP and α-tubulin antibody (in panel a) and with KIF2C and α-tubulin antibody (in panel b). DAPI was used to stain nucleus. (**B**) U-2OS cells were treated with control or KIF2C siRNA for 72 h and blots probed with KIF2C and α-tubulin antibody are shown. (**C**) U-2OS Cells were treated with control and KIF2C siRNA for 72 h and scored for binucleated cells. (**D, E**) Quantification of KIF2C intensity at centromere (see Figure 3**F**). Centromere intensity was measured from maximum intensity projection images using Zen software. Data represents MEAN ± S.D., ns: not significant (unpaired Student's t-test). Scale bars: 5 µm (**A**) and 20 µm (**C**).

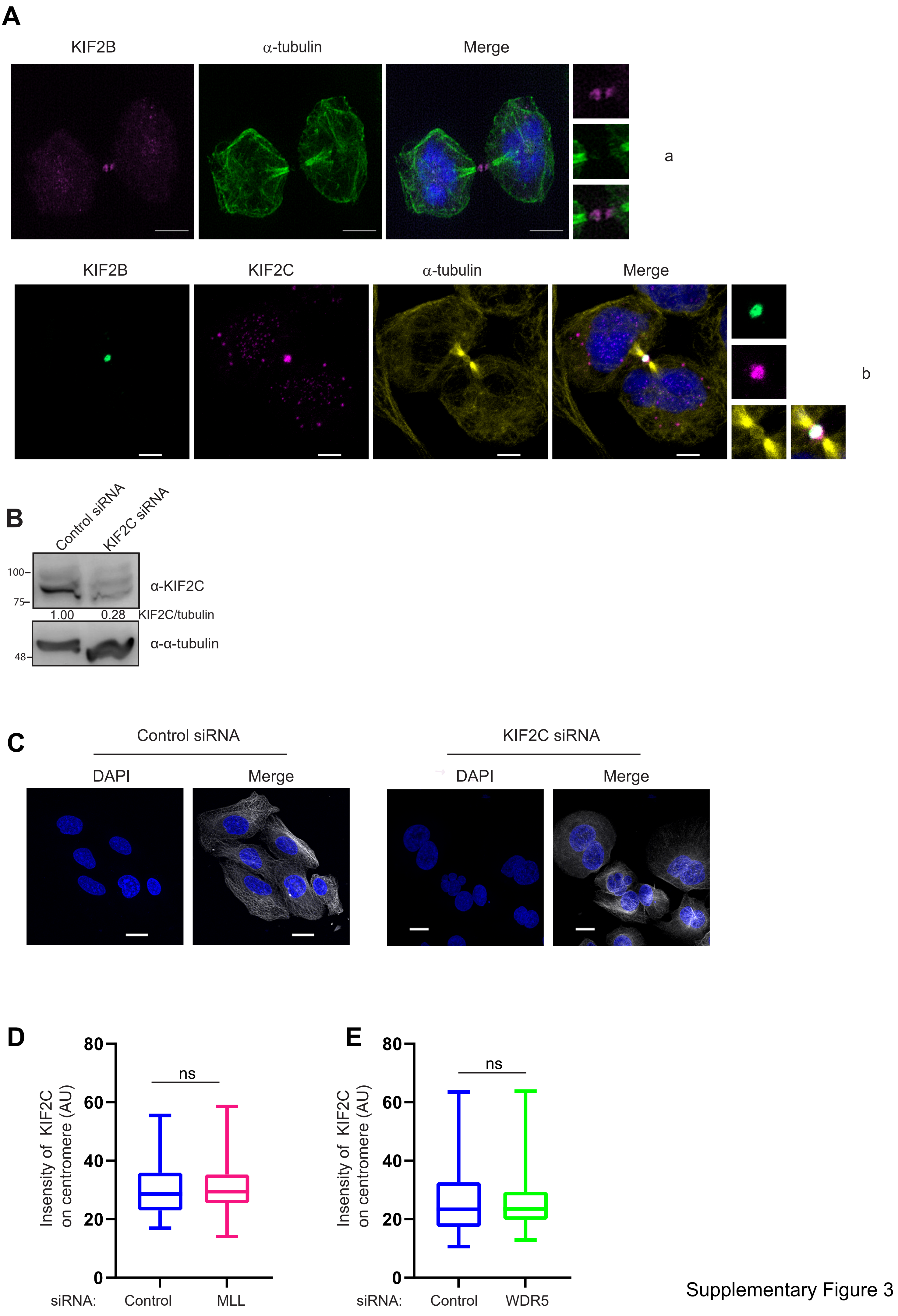


Supplementary Figure 3

**Figure S4**: **Localization of anillin during furrow ingression.** (**A**) Schematic of KIF2C full-length protein is shown. GFP-KIF2C wild type protein was modified using site directed mutagenesis to generate siRNA resistant construct with silent mutations as shown in the schematic. siRNA resistant construct was used to generate GFP-KIF2C G495A mutation. (**B**) U-2OS cells were fixed using paraformaldehyde and stained with Anillin and α-tubulin antibody. Image panels show localization of aniline as cell progresses from early to late anaphase and early to late telophase. White arrows indicate the cleavage furrow site. Scale bars: 5 μm

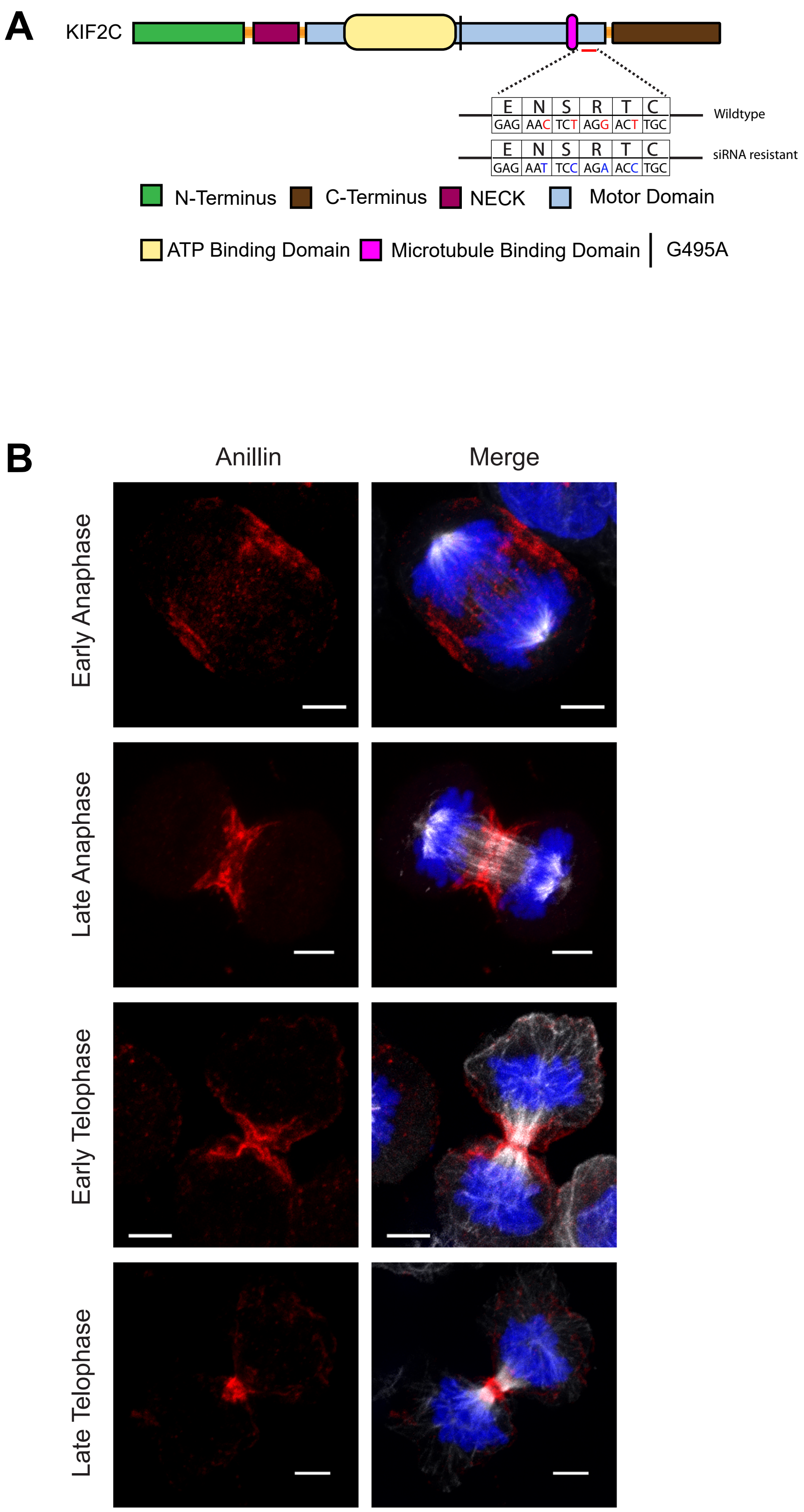


Supplementary Figure 4
